# Analysis of a Rpb2 mutant in *Schizosaccharomyces pombe* reveals a non-canonical role for Elp1 in regulating RNAi-dependent heterochromatin assembly

**DOI:** 10.1101/2025.07.02.662331

**Authors:** Mamta B. Nirmal, Maya E. Pearce, Cameron T. Liu, Jared M. Finkel, Katherine S. Darrow, Tommy V. Vo

## Abstract

Heterochromatin is a repressive epigenetic state that suppresses transcription and safeguards genomic integrity. However, the full mechanism of how it is regulated remains elusive. Here, we focus on a previously described Pol II variant called *rpb2-N44Y,* which has a single substitution mutation within the Rpb2 subunit of Pol II that reduces RNAi-dependent heterochromatin. Through CRISPR-mediated site-directed mutagenesis, we find that *rpb2-N44Y* is a gain-of-function mutation. Furthermore, the heterochromatin defects of the *rpb2-N44Y* mutant requires a subunit of the Elongator complex called Elongator Protein 1 (Elp1), a protein that canonically promotes in mcm^5^s^2^U_34_ tRNA modifications. Intriguingly, we find that loss of Elp1, but not of other Elongator subunits such as Elp3, can robustly suppress heterochromatin defects in the *rpb2-N44Y* mutant. Elp1 acts independently of the mcm^5^s^2^ U_34_ tRNA modification to suppress RNAi-dependent heterochromatin at the pericentromere and the levels of small interfering RNAs (siRNAs) at affected heterochromatin. Overall, our study reveals two distinct Rpb2-centric pathways, via RNAi or Elp1, that can positively or negatively regulate heterochromatin, respectively. Furthermore, our findings reveal the first evidence of a chromatin function for Elp1 that is distinct from its canonical role in tRNA modifications. This work expands our understanding of how Elp1 can influence chromatin biology.

**Article summary:** RNAi-dependent heterochromatin plays a key role in silencing gene expression from fission yeast to animals. However, it remains unclear what are all the factors that regulate this heterochromatin type. Here, the authors performed genetic interaction analyses to identify the conserved *elp1* gene as having the potential to regulate RNAi-dependent heterochromatin. Furthermore, the authors devised a separation-of-function mutant to find that this chromatin function of *elp1* is distinct from its canonical role in tRNA modifications. These findings expand our knowledge about the human disease-relevant *elp1* gene beyond its well-known roles in the Elongator complex and in tRNA modifications.

## Introduction

Eukaryotic chromatin consists of DNA and histone proteins that maintain the genetic information. The way in which chromatin is organized can determine how genetic information is used. Broadly, chromatin can adopt two distinct organizational states called euchromatin and heterochromatin. Euchromatin is a relaxed state that is accessible by transcription machinery and is often marked by transcription-associated histone modifications such as tri-methylation of histone H3 lysine-36 (H3K36me3) (Georgescu *et al*. 2020; Morrison and Thakur 2021). In contrast, heterochromatin is highly condensed and transcriptionally repressed. Molecular hallmarks of heterochromatin include the enrichment of di/tri-methylation of histone H3 lysine-9 (H3K9me2/3), H4K20me3, and histone hypoacetylation (Grewal and Jia 2007; Dambacher *et al*. 2013). The distribution of euchromatin and heterochromatin helps ensure overall genome integrity and proper gene regulation (Grunstein *et al*. 1995; Murakami 2013; Papamichos-Chronakis and Peterson 2013). Understanding how these states are assembled is important for determining how eukaryotes respond to developmental and environmental cues (Zhu *et al*. 2013).

The fission yeast *Schizosaccharomyces pombe* is a powerful model organism for studying heterochromatin because it utilizes conserved regulatory factors and mechanisms while exhibiting less genetic redundancy compared to higher eukaryotes and possessing high genetic tractability, thus facilitating the deduction of molecular pathways. As in animals and plants, heterochromatin in *S. pombe* is characterized by gene repression, position-effect variegation, and the presence of specific chromatin modifications such as H3K9me2/3 (Grewal and Jia 2007). H3K9 methylations are molecular platforms that recruit chromodomain proteins such as Swi6 (the yeast homolog of mammalian Heterochromatin Protein 1) and the histone deacetylase Clr3 (the yeast homolog of mammalian HDAC6/10) to mediate transcriptional repression and maintenance of the heterochromatic state (Sugiyama *et al*. 2007; Ragunathan *et al*. 2015; Seman *et al*. 2023). The initial establishment of H3K9me2/3 can occur via DNA- or RNA-dependent mechanisms. DNA-binding proteins such as Atf1 or Taz1 can recruit the sole H3K9 methyltransferase, Clr4, in *S. pombe* (Jia *et al*. 2004; Kanoh *et al*. 2005; Zofall *et al*. 2016). RNA-dependent H3K9me2/3 relies on long regulatory RNAs, such as meiotic transcripts containing Determinants of Selective Removal (DSR) sequences or small interfering RNAs (siRNAs) (Faber and Vo 2022). Mechanisms that couple H3K9me2/3 with siRNAs require RNA interference (RNAi) proteins, which are conserved between *S. pombe* and animals (Verdel *et al*. 2004; Shabalina and Koonin 2008; Andrews *et al*. 2014). This suggests a well-conserved role of the RNAi pathway in heterochromatin assembly across the eukaryotic tree of life.

In *S. pombe*, the establishment of RNAi-dependent heterochromatin initially requires Pol II-mediated transcription to generate precursor transcripts, which are then processed into mature siRNAs (Grewal and Elgin 2007; Holoch and Moazed 2015). Supporting the role of Pol II in heterochromatin assembly, previous studies have identified mutations that disrupt RNAi-mediated heterochromatin in three distinct Pol II subunits (Rpb1/2/7) (Djupedal *et al*. 2005; Kato *et al*. 2005; Inada *et al*. 2016; Kajitani *et al*. 2017). These mutants revealed that Pol II contributes to heterochromatin formation through mechanisms including RNA transcription, Mediator complex recruitment, RNA-chromatin interactions, and the recruitment of heterochromatin nucleation proteins. While most studies have focused on the Rpb1 subunit or proteins that interact with Rpb1 (Carlsten *et al*. 2012; Thorsen *et al*. 2012; Inada *et al*. 2016; Kajitani *et al*. 2017), the roles of other subunits remain less clear. Given that induced knockdown of Rpb2, but not Rpb1, in mouse embryonic stem cells reduces H3K9me3 (Bao *et al*. 2024), it is plausible that non-Rpb1 subunits of Pol II may uniquely regulate heterochromatin. This idea is further supported by a previously described mutant of Rpb2 in *S. pombe* called *m203,* which remains transcriptionally competent but disrupts RNAi-dependent heterochromatin (Kato *et al*. 2005). This phenotype is distinct from previously characterized Rpb1 and Rpb7 mutants, which are altered in both transcriptional and heterochromatin-forming potentials (Djupedal *et al*. 2005; Kajitani *et al*. 2017). The *m203* variant could be a useful genetic tool to reveal new insights into how heterochromatin formation occurs downstream of the initial transcription step.

Here, we show that *m203* is a gain-of-function genetic mutation that is a useful tool to identify novel regulators of RNAi-dependent heterochromatin. Through a genetic suppressor screen of *m203* mutant cells, we found that the Elp1 protein can suppress the accumulation of siRNAs at an RNAi-dependent heterochromatic reporter locus. Although Elp1 is primarily known as a scaffolding subunit of the Elongator complex, which promotes 5-methoxycharbonylmethyl-2-thiouridine (mcm^5^s^2^) modification of tRNAs (Huang *et al*. 2005; Dauden *et al*. 2019), we found that neither mutations of other Elongator complex-associated factors nor global inhibition of translation could mimic the pronounced impact of Elp1 loss on RNAi-dependent heterochromatin. Furthermore, genetic inhibition of Elp1-dependent mcm^5^s^2^ tRNA modification failed to restore heterochromatic silencing in *m203* cells. This suggests that Elp1 could have a moonlighting role in epigenetic gene regulation. To our knowledge, this work demonstrates the first case where Elp1 has a role that is largely independent of the Elongator complex and the mcm^5^s^2^U_34_ tRNA modification. This may help explain why the Elp1 homolog in humans is associated with distinct disease pathologies compared to the other subunits of the Elongator complex (Gaik *et al*. 2023).

## Materials and Methods

### Yeast culturing and manipulation

*S. pombe* yeast strains that were used in this study are indicated in Supplementary Table 1. Strains were generated by using standard genetic crossing, by PCR-based gene deletion with transformation by lithium acetate method (Krawchuk and Wahls 1999), or SpEDIT CRISPR-Cas9 editing (Torres-Garcia *et al*. 2020). All yeast were cultured in medias that were based on the yeast extract rich medium with adenine supplement (YEA) and incubated at 32°C for 2-4 days. Liquid cultures were grown at 32°C with 220 rpm shaking for aeration. As appropriate for our experiments, we have used YEA media with the following: 5-FOA (GoldBio, 850 mg/L), cycloheximide (Sigma, 20 mg/mL), G418 sulfate (GoldBio, 50 mg/L), hygromycin (GoldBio, 200 µg/mL), and nourseothricin (GoldBio, 100 mg/L).

### Serial Spotting Assays

Yeast cells were pre-cultured at 32°C for 220 rpm overnight until cells reached log growth phase. Next, cell optical densities (OD_600_) were measured using a Denovix DS-11+ apparatus and cells were normalized to the equivalent OD_600_ of 0.2 – 0.5. Finally, cells were serially diluted 4-fold and equal volumes of cells per spotted onto the indicated media plates. Spotted plates were incubated at 32°C without shaking for 2-4 days. Images were collected using a BioRad Chemidoc imager using the Colorimeteric setting. Any image correction was uniformly applied to the entirety of the shown images.

### Generation of tRNA overexpression plasmids

tRNA expression plasmids were generated similar to how it was described in (FernÁndez-VÁzquez *et al*. 2013) with the major modification being that the plasmids were designed to be selectable with nourseothricin (Nat). This allowed us to culture our cells uniformly in YEA-based media, minimizing phenotypic variation due to major differences in media composition. Briefly, the pRep41X plasmid (Addgene) was modified by replacing the *LEU2* gene with the *natMX* resistance gene between the HindIII and NheI restriction sites. The resulting construct was designated pRep_Nat. Next, the *SPBTRNALYS.06* (encoding for tRNA^Lys^_UUU_) and *SPCTRNALYS.11* (encoding for tRNA^Lys^_CUU_ genes, along with 500 bp upstream and downstream DNA sequences, were PCR-amplified from the *S. pombe* genome as previously described by (FernÁndez-VÁzquez *et al*. 2013). These PCR products were cloned into the NheI site of the pRep_Nat plasmid. Plasmids used in this study are listed in Supplementary Table 2.

### Derived Cleaved Amplified Polymorphic Sequences (dCAPS)

Yeast cells were first grown on YEA-based media plate. Next, cells were treated with zymolyase (Zymo Research) at 30°C for 45 minutes and lysed by heating to 95°C for 10 minutes. The yeast lysate, containing the released genomic DNA, was used as template for PCR (with GoTaq DNA polymerase) to amplify part of the *rpb2* gene to include the codon that encodes for the 44^th^ amino acid residue. PCR oligos are listed in Supplementary Table 3. Next, the PCR products were directly subjected to restriction digestion with BstZ171-HF (NEB). Finally, an aliquot of digested PCR products were resolved in a 1% agarose gel that was pre-stained with ethidium bromide.

### Chromatin Immunoprecipitation (ChIP)–qPCR

For all ChIP experiments, *S. pombe* cells were inoculated into 50 mL of YEA medium and incubated overnight at 32°C with shaking at 220 rpm to an OD600 of 0.5–0.6. Cells were crosslinked with 1% formaldehyde for 20 minutes at room temperature, followed by quenching with 2.5M glycine. Then, cell pellets were washed twice with 20 mL of cold 1× PBS. The cell pellet was resuspended in 350 µL of ChIP lysis buffer (50 mM HEPES pH 7.5, 140 mM NaCl, 1 mM EDTA, 1% Triton X-100, 0.1% sodium deoxycholate, EDTA free 1X cOmplete) and disrupted with 1 mL of glass beads using bead-beating (1 min beating, 2 min on ice, repeated 3 times). Chromatin was sheared using a QSonica Q800R sonicator (20 sec ON / 40 sec OFF cycles, 70% amplitude) for 10 minutes at 4°C. Cell debris was removed by centrifugation at 4°C, 1,500×g for 5 minutes. The supernatant was transferred to fresh 1.5 mL tubes and diluted with ChIP lysis buffer. Lysates were precleared with 20 µL of protein A/G agarose beads (Santa Cruz, cat no. A3124) and incubated at 4°C for 1 hour. 50 µL of pre-cleared lysate was reserved as the 5% input control. The remaining lysate was incubated with the H3K9me3 (Active Motif, cat no. 39062) antibody and protein A/G agarose beads for overnight at 4°C. Beads were washed sequentially (twice each) with the following buffers: ChIP Buffer I (50 mM HEPES pH 7.5, 140 mM NaCl, 1 mM EDTA, 1% Triton X-100, 0.1% deoxycholate), ChIP Buffer II (same as Buffer I, but with 500 mM NaCl), ChIP Buffer III (10 mM Tris-HCl pH 8.0, 250 mM LiCl, 0.5% NP-40, 0.5% deoxycholate, 1 mM EDTA), and 1x Tris-EDTA buffer.

Immunoprecipitated chromatin was extracted by incubating the agarose beads with 100 µL of elution buffer (50 mM Tris-HCl pH 8.0, 10 mM EDTA) at 65 °C and agitating at 1,000 rpm for 30 minutes. This step was repeated to obtain a total volume of 200 µL eluant. To each ChIP sample, 4 µL of 5N NaCl and 1µL RNase A (Thermo Fisher Scientific, 20 mg/mL) was added, and samples were incubated overnight at 65°C in a water bath to reverse crosslinks. Similarly, input samples were adjusted to 200 µL with elution buffer, followed by the addition of 2.6 µL of 5N NaCl and 1 µL RNase A and incubated overnight at 65°C in a water bath to reverse crosslinks. After sample tubes were cooled to room temperature, 1µL of Proteinase K (Thermo Fisher Scientific) was added to each tube and incubated at 50°C for 1 hour. DNA was purified by ethanol precipitation using 3 M sodium acetate, 1 µL glycogen and resuspended into 50 µL of molecular-grade water. For qPCR, 1 µL of ChIP DNA was used in each reaction with iTaq SYBR Green reagents (BioRad). PCR oligos used for qPCRs are listed in Supplementary Table 3. Antibodies used for ChIP are listed in Supplementary Table 4.

### Western Blot

For protein extraction, 2 OD_600_ units of yeast cells were harvested by centrifugation into 2 mL screw-cap tubes. The cell pellet was resuspended in 200 µL of cold 20% trichloroacetic acid (TCA). Approximately 400 µL of acid-washed glass beads were added, and the cells were lysed using mechanical disruption with a bead beater (20 seconds beating × 3 cycles with 45-second intervals on ice). Following lysis, each tube was punctured, placed on 5ml polystyrene collection tube and the lysate was collected by centrifugation at 1000 rpm for 30 seconds. The remaining beads were washed with 400 µL of cold 5% TCA, spun again, and the wash was pooled with the initial lysate. The combined lysate (∼600 µL total) was transferred to a 1.5 mL microcentrifuge tube and centrifuged at 13,000 rpm for 5 minutes at 4°C to pellet the proteins. Supernatant was discarded and centrifuged 2-3 times to remove residual TCA. The resulting protein pellet was resuspended in 50–100 µL of SDS-PAGE 1x sample loading buffer (Thermo Fisher Scientific, cat no. NP0007-2988243). Pellets were often difficult to dissolve and required heating to fully solubilize. Samples were then boiled at 95°C and loaded on SDS-PAGE 4–12% Bis-Tris gels (Thermo Fisher Scientific, cat no. 25010870). Electrophoresis was performed in an Invitrogen mini tank apparatus using NuPage 1X running buffer at room temperature at 200 volts for 45 minutes. Proteins were transferred onto PVDF membranes (pre-activated with methanol) using wet transfer (Nupage 1X transfer buffer (NP0006-2988243) at 20 volts for 60–90 minutes at room temperature.

Following the transfer, membranes were blocked for 1 hour at room temperature in 1X TBST (Tris-buffered saline with 0.1% Tween-20) containing 5% non-fat dry milk. Membranes were then incubated overnight at 4°C with 1:2,000 mouse primary antibody that was diluted in blocking buffer. After washing three times with 1X TBST (10 minutes each), membranes were incubated with an HRP-conjugated secondary anti-mouse (Jackson laboratory, cat no. 115-035-146) for 1 hour at room temperature. Following three additional 1X TBST washes, signal detection was performed using an enhanced chemiluminescence (Cytiva, cat no. RPN2232-18206643) substrate and visualized using the BioRad chemiluminescence imaging system. Antibodies used are listed in Supplementary Table 4.

### Isolation of total RNA

Total RNA was isolated from 10mL of *S. pombe* cells that were grown in liquid YEA media at 32°C to OD_600_ between 0.5 – 1. Before starting the isolation procedure, all centrifuges were set to 6°C, and 100% isopropanol was kept at –20°C. A water bath was set to 65°C with enough water to submerge half of a 15 mL Falcon tube. Heavy phase lock tubes (Qiagen) were centrifuged and placed on ice. The cell pellet was thawed on ice (if frozen), and any excess media was removed. 1 mL of AES buffer (50mM sodium acetate pH 5.3, 10mM EDTA, 1% SDS) was added to the cell pellet and mixed gently using pipetting. In a fume hood, 1 mL of acidic phenol (pH 4.5) was added to the cell suspension and gently mixed by inverting the tube 10 times. Next, the sample was vortexed for 10 seconds and incubated in the 65°C water bath for 45 minutes. This incubation was repeated five times without interruption. The resulting cell-AES-phenol mixture was transferred into heavy phase lock tubes and centrifuged at 4,000g for 5 minutes at 6°C, taking care not to disturb the two layers. Next, 1 mL of phenol:chloroform:isoamyl alcohol (25:24:1) (VWR) was added to the sample, and the tube was inverted gently 10 times to mix, followed by centrifugation at 4,000g for 5 minutes at 6°C. In the fume hood, 1 mL of chloroform was added to each sample, gently mixed by inversion (10 times), and centrifuged again at 4,000g for 4 minutes at 6°C. Carefully, 700 µL of the top clear aqueous phase was pipetted into a 1.5 mL microcentrifuge tube. To this, 70 µL of 3M NaOAc and 1 µL of glycogen were added and mixed thoroughly by inverting the tube 10 times. Ice-cold 100% isopropanol was then added, and the tube was inverted 10 times to mix. The sample was centrifuged at maximum speed for 20 minutes at 6°C. A moderate RNA pellet was observed, the supernatant was discarded, and the pellet was washed with 900 µL of 75% ethanol by inverting the tube 10 times, followed by centrifugation at maximum speed for 5 minutes at 6°C. The supernatant was removed using a P1000 pipette, and any residual liquid was removed using a P200 or P20 pipette. The pellet was air-dried for 10 minutes before adding 100 µL of DEPC-treated water. The sample was incubated at room temperature for 5 minutes to dissolve the pellet and mixed gently by pipetting. RNA concentration was measured using a DS-11+ spectrophotometer. Then, 10 µg of isolated RNA was then treated with Turbo DNase (Thermo Fisher Scientific, cat no. AM2238) in 50ul of total volume to remove genomic DNA. Then, the volume was brought to 100ul by adding DEPC water. 100 µL of phenol:chloroform:isopropyl alcohol was added. The sample was gently inverted 10 times to mix and centrifuged at 20,000 x g for 5 minutes at 6°C to separate the organic (bottom) and aqueous (top) layers. 70 µL of the top aqueous layer was carefully transferred to a fresh microcentrifuge tube. To this, 7 µL of 3M sodium acetate and 1 µL of glycogen were added and mixed by gentle tapping. Then, 235 µL of 200-proof ethanol was added and mixed by inverting the tube 40 times. The tubes were incubated overnight at –20°C to precipitate the RNA. The sample was centrifuged at maximum speed (20,000 x g) for 30 minutes at 6°C to pellet the purified RNA. After centrifugation, the supernatant was carefully discarded using a P1000 pipette, leaving behind the RNA pellet. To wash the pellet, 900 µL of 75% ethanol was added, and the tube was inverted 10 times to mix. The sample was then centrifuged again at 20,000 x g for 5 minutes at 6°C. As much of the supernatant as possible was removed, followed by a brief spin to collect residual liquid, which was removed using a P10 pipette. The pellet was air-dried for 5 minutes to allow the remaining ethanol to evaporate. Finally, 20 µL of DEPC-treated water was added to dissolve the RNA. The tube was incubated at room temperature for 10 minutes, mixed gently by pipetting, and the RNA concentration was measured. Successful DNase treatment was validated by synthesizing cDNA using the Invitrogen SuperScript III Reverse Transcriptase kit, followed by PCR to confirm the absence of genomic DNA contamination. Tapestation analysis at the MSU Genomics core facility was performed to confirm the quality of RNA based on RIN values. Then the samples were submitted to ArrayStar MS in 1/10 volume of 3M sodium acetate and 350 μL of 100% ethanol.

### Small RNA sequencing

Total RNA (free from contaminating genomic DNA) was used to prepare small RNA libraries with 18-40 bp inserts. Briefly, 3’ and 5’ Illumina sequencing adapters were ligated to the end of RNAs. Next, cDNA was generated and subsequently used for PCR amplification. Next, libraries were resolved using gel electrophoresis and were size-selected. Sequencing of the small RNA-seq libraries was performed on an Illumina NovaSeq with single-end reads of 50 nt. Approximately 10-14 million raw reads were generated per sequenced sample.

The raw sequencing data was first assessed by FastQC to ensure high-quality data. Next, Cutadapt was used to remove poor quality reads, reads with lengths below 15 nt, and adapter sequences. The remaining reads (74-80% of total raw reads) were aligned to the *S. pombe* genome (May 2012 version) using Bowtie. Samtools was used to keep alignments from reads of lengths 21-24 nt. After, only uniquely mapped reads (-q 40) were used for coverage calculations. Coverage was calculated using the per million scaling factor. Additionally, total coverage was determined as the sum of coverage on the positive strand (having positive coverage) and on the negative strand (having negative coverage). siRNAs that are generated through bidirectional transcription and/or the activity of RNA-dependent RNA polymerase typically show total coverage that spans both positive and negative strands (Yamanaka *et al*. 2013).

### Detection of tRNA modifications by liquid chromatography mass spectrometry

Transfer RNAs (tRNAs) were isolated from total RNA (free from contaminating genomic DNA) using urea-PAGE electrophoresis to extract 70-90nt small RNAs. Once isolated, the tRNAs underwent enzymatic digestion to produce dephosphorylated, single nucleosides. These purified nucleosides were subsequently analyzed by LC-MS/MS to identify modified nucleosides. Signal intensities were normalized by the amount of total tRNAs that were loaded. Finally, the resulting data were processed and normalized using Agilent Qualitative Analysis software, where multiple reaction monitoring (MRM) peaks for each modified nucleoside were extracted.

## Results

### Gain-of-function Rpb2 mutations promote heterochromatic defects

The *m203* mutant is defined by an asparagine (Asn) to tyrosine (Tyr) substitution at amino acid residue 44 of Rpb2 (N44Y), the second largest subunit of Pol II (Kato *et al*. 2005; SpÅhr *et al*. 2009). Cells of this genotype lose heterochromatic silencing of a *ura4^+^* reporter gene, which has been inserted into the pericentromere of chromosome 1 (*otr1R::ura4^+^*) (Kato *et al*. 2005) (Fig. 1a). While genetic analyses have indicated that *m203* leads to heterochromatin defects through the disruption of the RNAi pathway (Kato *et al*. 2005), it remains unclear how this occurs. We first determined whether *m203* is a loss- or gain-of-function mutation by performing CRISPR-Cas9 site-directed mutagenesis to substitute Asn^44^ with other amino acid residues and measuring their impacts on the repression of the *otr1R::ura4^+^* reporter (Fig. 1a). In wild-type cells with Rpb2-Asn^44^ (*WT*), heterochromatin forms at the *otr1R::ura4^+^* reporter which leads to transcriptional repression of *ura4^+^* (Allshire *et al*. 1994). Accordingly, wild-type cells can readily form yeast colonies in the presence of the drug 5-fluoroorotic acid (5-FOA) (Fig. 1b). Cells that do not express *ura4^+^* will not convert 5-FOA into a toxic metabolite and will, therefore, be able to proliferate (Verdel *et al*. 2004; Kato *et al*. 2005; Cam and Whitehall 2016). For mutant cells that fail to grow in the presence of 5-FOA, such as the case for *m203* cells (Kato *et al*. 2005), they could have lost heterochromatic repression of the *otr1R::ura4*^+^ reporter, and thus 5-FOA is metabolized into a toxic product that inhibits growth (Fig. 1b). We tested the hypothesis that the similarity of amino acid side-chains would predict the growth patterns of cells on media with 5-FOA. Cells in which Rpb2-Asn^44^ was substituted with glutamine (Glu, Q), whose side-chain is most similar to that of Asn, grew similarly to wild-type in the presence of 5-FOA (Fig. 1b). Vice versa, cells where Rpb2-Asn^44^ was substituted with phenylalanine (Phe, F), which has a bulky side-chain similar to Tyr, grew as poorly as *rpb2-N44Y* (*m203*) cells on media with 5-FOA (Fig. 1b). We further validated by chromatin immunoprecipitation with quantitative PCR (ChIP-qPCR) that both *rpb2-N44Y* and *rpb2-N44F* mutants decreased the levels of heterochromatic H3K9me3 at the *otr1R::ura4^+^* reporter (Fig. 1c). Together, our results support the notion that the side-chain of the 44^th^ amino acid of Rpb2 is important for pericentromeric heterochromatin.

**Fig 1.**
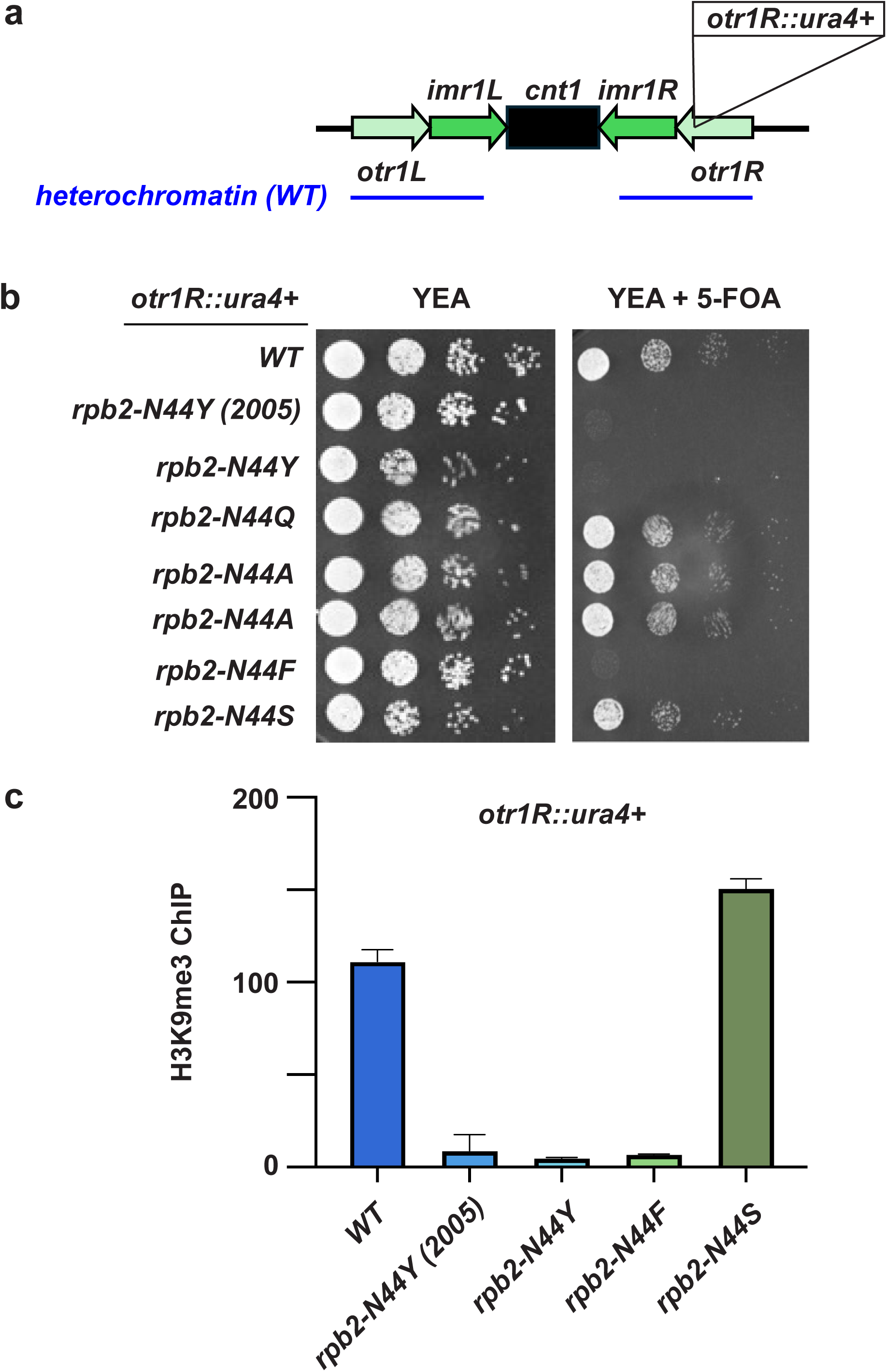
The 44^th^ amino acid residue of Rpb2 is a critical determinant of heterochromatin status at the pericentromere. (A) Schematic illustration of the pericentromeric *otr1R::ura4^+^* reporter that is used in this work. (B) Serial spotting assay to assess the impact of *rpb2* substitution mutations on yeast growth on media −/+ 5-FOA. The wild-type strain is denoted by *WT*. The *rpb2-N44Y* (2005) yeast strain was previously used in (Kato *et al*. 2005). The other *rpb2-N44Y* yeast strain was generated by our lab using CRISPR-Cas9. (C) ChIP enrichment of H3K9me3 at *otr1R::ura4^+^* in yeast strains of various *rpb2* genotypes. Enrichments were normalized relative to the euchromatic *leu1* gene and a 5% input control. Error bars indicate standard deviation based on *n* ≥ 2.

To test whether *rpb2-N44Y* and *rpb2-N44F* are loss- or gain-of-function mutations, we substituted Rpb2-Asn^44^ with alanine or serine (*rpb2-N44A* or *rpb2-N44S*), which are amino acids that are highly dissimilar with the other amino acids that were tested. We found that cells with *rpb2-N44A* and *rpb2-N44S* variants proliferated similarly to wild-type cells in the presence of 5-FOA (Fig. 1b). Furthermore, there was still H3K9me3 enrichment at *otr1R::ura4^+^* in the *rpb2-N44S* variant (Fig. 1c). This suggests that the dissimilar variants *rpb2-N44*, *-N44A*, and *-N44S* can all support heterochromatic repression of the *otr1R::ura4^+^* reporter. We propose that *rpb2-N44Y* and *rpb2-N44F* are gain-of-function mutations that can disrupt the assembly of H3K9me3-marked heterochromatin. Supporting this conclusion, substitution of N44 with tryptophan, lysine, methionine, or glutamic acid resulted in wild-type growth phenotypes on media with 5-FOA (Fig. S1), indicating that only specific substitutions at the 44^th^ position of Rpb2 lead to loss of heterochromatic repression.

### Heterochromatin loss in *rpb2-N44Y* and *-N44F* mutants require the *elp1* gene

To identify the factors that mediate heterochromatin loss in the *rpb2-N44Y* (in *m203* cells) and *rpb2-N44F* mutant cells, we sought suppressor mutations that could restore the repression of our *otr1R::ura4^+^*reporter. To achieve this, we first deleted individual genes that are associated with transcription elongation to test whether any single deletion could restore heterochromatic repression of our *otr1R::ura4^+^* reporter in *m203* cells (Fig. 2a). We deleted the *tfs1* gene (encodes for a homolog of mammalian TFIIS) because this mutant suppresses heterochromatic defects in cells that lack the RNAi gene called *ago1* (Reyes-Turcu *et al*. 2011). We also deleted the *rpb9* and *elp1* genes because, similar to *tfs1*, they have been implicated in the regulation of transcription elongation (Otero *et al*. 1999; Hemming *et al*. 2000). We discovered that loss of *elp1* (*elp1*Δ) in *m203* cells enabled strong growth in the presence of 5-FOA, similar to wild-type cells (Figs. 2a and S2a). The restored growth phenotype of *m203 elp1*Δ cells on 5-FOA media persisted through genetic crosses, indicating that the growth phenotypes are dependent on the *rpb2* and *elp1* genotypes (Fig. 2b). Furthermore, ChIP-qPCR confirmed that *elp1*Δ increased heterochromatic H3K9me3 and decreased euchromatic H3K36me3 at our reporter locus in *m203* cells (Figs. 2c, d). The changes in H3K9me3 likely affect the expression state of our *otr1R::ura4^+^*reporter because *m203 elp1*Δ cells reverted to hypersensitivity on 5-FOA-containing media after the loss of the *clr4* gene (encodes for the sole H3K9 methyltransferase) (Fig. 2e). Lastly, we independently confirmed that *elp1*Δ in *rpb2-N44F* mutant cells also restored robust cell growth on 5-FOA media (Figs. S2a,b). Collectively, our results suggest that the *rpb2-N44Y* and *rpb2-N44F* mutants promote an *elp1*-dependent process, which reduces the heterochromatic repression of our pericentromeric reporter.

**Fig 2.**
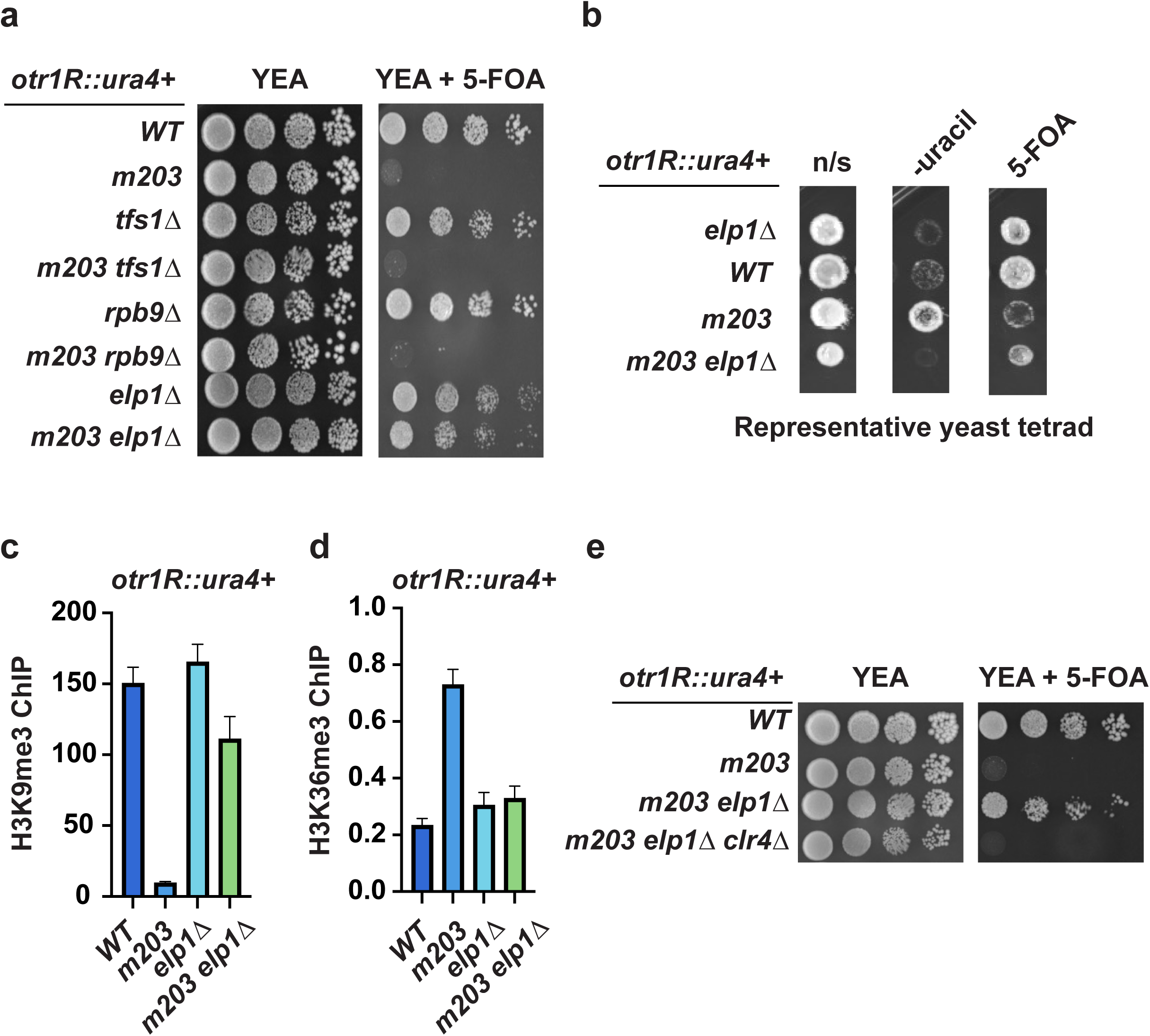
Loss of elp1 restores heterochromatic repression of the *otr1R::ura4^+^* reporter in *m203* (*rpb2-N44Y*) cells. (A) Serial spotting assay to determine gene deletions that can restore 5-FOA resistance in *m203* cells. (B) Tetrad analysis reveals the segregation of the 5-FOA resistance phenotype follows the *elp1*Δ genotype. Shown is the growth phenotypes for yeast derived from a representative tetrad. Cells were grown on non-selective (n/s) YEA media, media lacking uracil, or media containing 5-FOA. (C) ChIP enrichment of heterochromatic H3K9me3 at *otr1R::ura4^+^* in the indicated yeast strains. (D) ChIP enrichment of euchromatic H3K36me3 at *otr1R::ura4^+^* in the indicated yeast strains. (E) Spotting assay showing that loss of the H3K9 methyltransferase *clr4* reverts the 5-FOA sensitivity of *m203 elp1*Δ cells to mirror the sensitivity of *m203* cells. ChIP enrichments were normalized relative to the euchromatic *leu1* gene and a 5% input control. Error bars indicate standard deviation and experiments were based on *n* ≥ 2.

We considered the possibility that Elp1 could regulate the *ura4^+^*gene outside of the heterochromatin context. To test this, we examined the impact of *elp1* loss on cells that expressed *ura4^+^* from its native euchromatic locus. Similar to wild-type cells with *ura4^+^* in euchromatin, *elp1*Δ cells robustly grew on media lacking uracil but not on media containing 5-FOA (Fig. S2c). This suggests that loss of *elp1* does not intrinsically alter *ura4* expression. Therefore, *elp1*-dependent regulation of *otr1R::ura4^+^* expression is likely based on the chromatin state.

Loss of the *elp1* gene enhances DNA damage sensitivity (Li *et al*. 2009; Chen *et al*. 2021). Therefore, we tested the possibility that *elp1*Δ could revert the *rpb2-N44Y* mutation of *m203* cells back to the wild-type allele (*rpb2-N44*) to restore heterochromatin. Towards this end, we isolated genomic DNA from *m203 elp1*Δ double mutant cells and performed Derived Cleaved Amplified Polymorphic Sequences (dCAPS) (Kaundun and Windass 2006) to probe for the wild-type and *m203* alleles of *rpb2*. We detected *rpb2-N44Y* as the dominant allele in *m203 elp1*Δ cells (Fig. S2d), suggesting that *elp1* loss does not restore repressive heterochromatin by majorly promoting the correction of the *rpb2* mutant allele. Rather, there is a positive genetic interaction between *elp1* and *rpb2-N44Y* that influences the chromatin state of our *otr1R::ura4+* reporter.

### Elp1 can also regulate some native heterochromatin domains

Outside of our pericentromeric reporter, we found that *m203* reduced di-methylation of H3K9 (H3K9me2) at a native heterochromatin region known as *HOOD-23*, which encompasses the *pho1* gene (Fig. 3a) (Yamanaka *et al*. 2013). At this region, H3K9me2-marked heterochromatin assembles upon the loss of the RNA exosome subunit Rrp6 (Yamanaka *et al*. 2013; Shah *et al*. 2014). Here, we found that *elp1* loss restored H3K9me2/3 in *m203* cells. We also tested whether *elp1*Δ can restore H3K9me2 at another native heterochromatin region known as *HOOD-10* (which encompasses the Tf2 retrotransposon element *tf2-5*) (Yamanaka *et al*. 2013). However, we did not find H3K9me2 enhancement in *m203 rrp6*Δ *elp1*Δ cells (Fig. 3b). This suggests that the role of Elp1 varies depending on the heterochromatin region. Overall, our results suggest that Elp1 could negatively regulate heterochromatin formation at diverse native and reporter loci.

**Fig 3.**
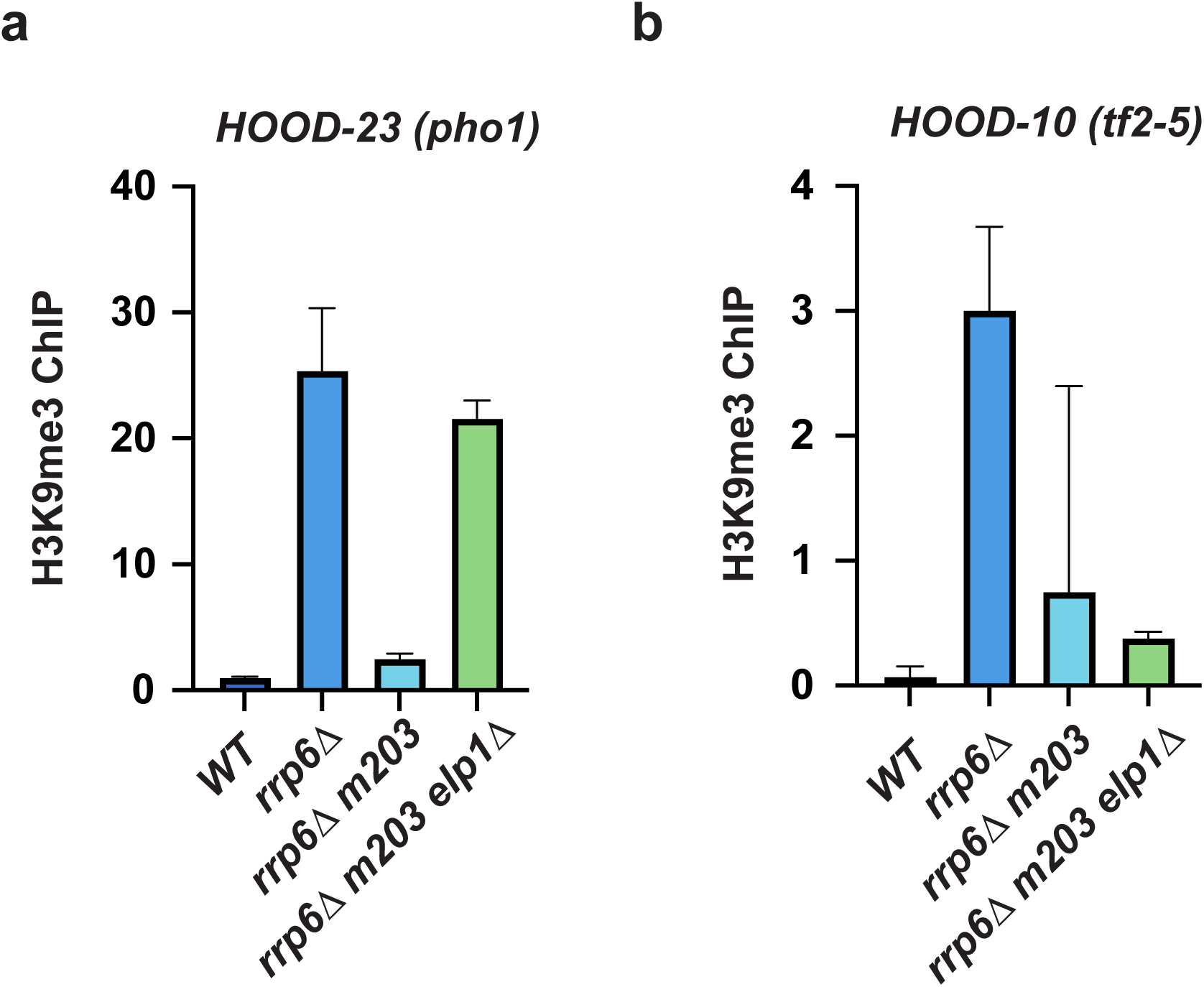
Loss of *elp1* restores heterochromatic H3K9me2 at certain heterochromatin domains. (A) ChIP enrichment of heterochromatic H3K9me2 at *HOOD-23*, which encompasses the annotated *pho1* gene, in the indicated yeast strains. (B) ChIP enrichment of heterochromatic H3K9me2 at *HOOD-10*, which encompasses the annotated *tf2-5* gene, in the indicated yeast strains. At both HOODs, H3K9me2 is known to be induced upon the loss of the *rrp6*, which encodes for a catalytic subunit of the nuclear exosome. ChIP enrichments were normalized relative to the euchromatic *leu1* gene and a 5% input control. Error bars indicate standard deviation based on *n* ≥ 2.

### Elp1 can suppress heterochromatic RNAi

The RNAi pathway is critically important for promoting heterochromatic repression of the *otr1R::ura4+* reporter (Verdel *et al*. 2004), prompting us to investigate whether Elp1 could act via that pathway. First, we performed small RNA sequencing and mapped 21 – 24 nucleotide (nt) siRNAs to the *S. pombe* genome. We analyzed levels of siRNAs that are derived from pericentromeric *dh* repeats because they promote H3K9me3-marked heterochromatin at our reporter (Verdel *et al*. 2004). We found that *elp1*Δ partially increases levels of *dh*-derived siRNA in *m203* cells (Fig. 4a). Next, we tested whether repression of our *otr1R::ura4^+^*reporter in *m203 elp1*Δ cells required the RNAi pathway by deleting *ago1* or *dcr1* in this strain to generate triple mutant strains *m203 elp1*Δ *ago1*Δ or *m203 elp1*Δ *dcr1*Δ (Figs. 4b). The *ago1* and *dcr1* genes encode for homologs of metazoan RNAi factors Argonaute or Dicer proteins, respectively (Tabara *et al*. 1999; Zamore *et al*. 2000; Bernstein *et al*. 2001; Verdel *et al*. 2004). We performed spotting assays in the absence or presence of 5-FOA, as previously described (Fig. 1b). In both cases, the absence of *ago1* or *dcr1* reverted cells back to the 5-FOA-sensitive state, suggesting that *elp1*Δ depends on the RNAi pathway to repress the *otr1R::ura4^+^* reporter (Figs. 4b). In *elp1*Δ cells with wild-type *rpb2* gene, RNAi-dependent repression dominants the *otr1R::ura4^+^* reporter and knockout of *elp1* has no apparent effect (Fig. S3). This suggests that *rpb2* could regulate the contribution of Elp1 and RNAi towards negative and positive regulation of heterochromatin, respectively.

**Fig 4.**
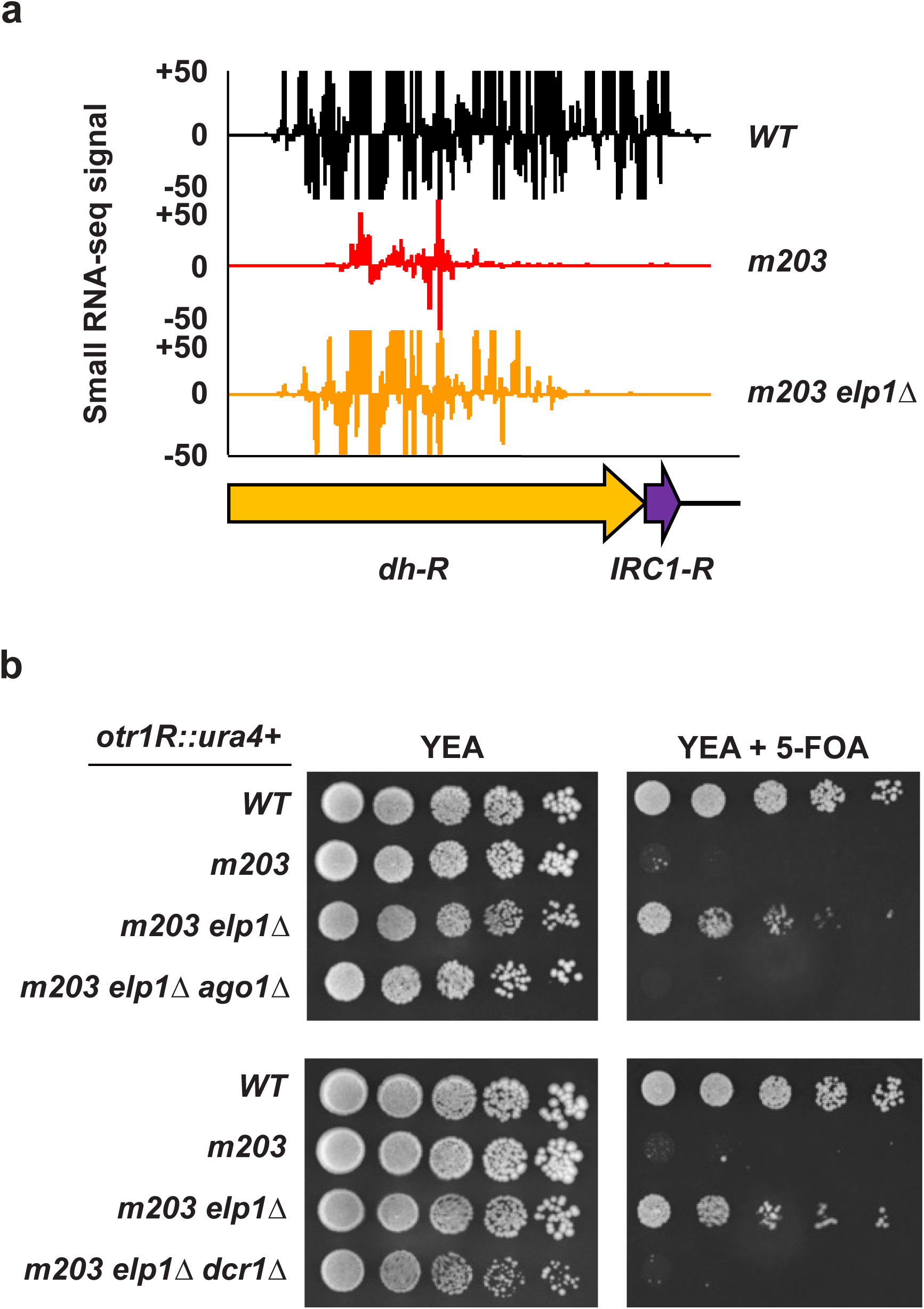
Heterochromatin regulation by *elp1* requires RNA interference. (A) Coverage of mapped 21 – 24 nucleotide Illumina reads obtained from small RNA-seq for the indicated yeast strains at the native *dh* repeats within the *otr1R* pericentromeric locus. The coverage was calculated as the sum across both positive and negative strands. (B) Spotting assays showing that loss of the RNAi-related genes *ago1* or *dcr1* enhances 5-FOA sensitivity in *m203 elp1*Δ cells.

### The Elp1 subunit of the Elongator complex distinctly contributes to heterochromatin

We next investigated how Elp1 could regulate RNAi-mediated heterochromatin. It is known that Elp1 is a core subunit of the Elongator complex, which is comprised of six proteins (Elp1-6) (Otero *et al*. 1999; Winkler *et al*. 2001; Jaciuk *et al*. 2023). Within this complex, Elp1 has a scaffolding function by interacting with the other subunits, and it also binds tRNA substrates to promote Elongator complex-mediated tRNA modifications (Dauden *et al*. 2019; Jaciuk *et al*. 2023). We tested whether ablating the other core subunits of the Elongator complex in *m203* cells would phenocopy the deletion of *elp1* and restore heterochromatin (Fig. 2a). To do this, we knocked out the *elp3* or *elp5* genes (*elp3*Δ or *elp5*Δ) in *m203* cells that carried our *otr1R::ura4^+^* reporter. While *m203* cells that lacked Elp1 (*m203 elp1*Δ) robustly grew on media with 5-FOA (Fig. 2a), *m203* cells lacking *elp3* or *elp5* only had modest growths on the same 5-FOA containing media (Fig. 5a). These phenotypes correlated with partial increases in H3K9me3 at the *otr1R::ura4+* reporter in *m203* cells lacking *elp3*, compared to a higher H3K9me3 restoration in *m203 elp1*Δ cells (Fig. 5b), compared to *m203* cells. Our results suggest that *m203* is more dependent on Elp1 than the other Elongator complex subunits to reduce heterochromatic repression.

**Fig 5.**
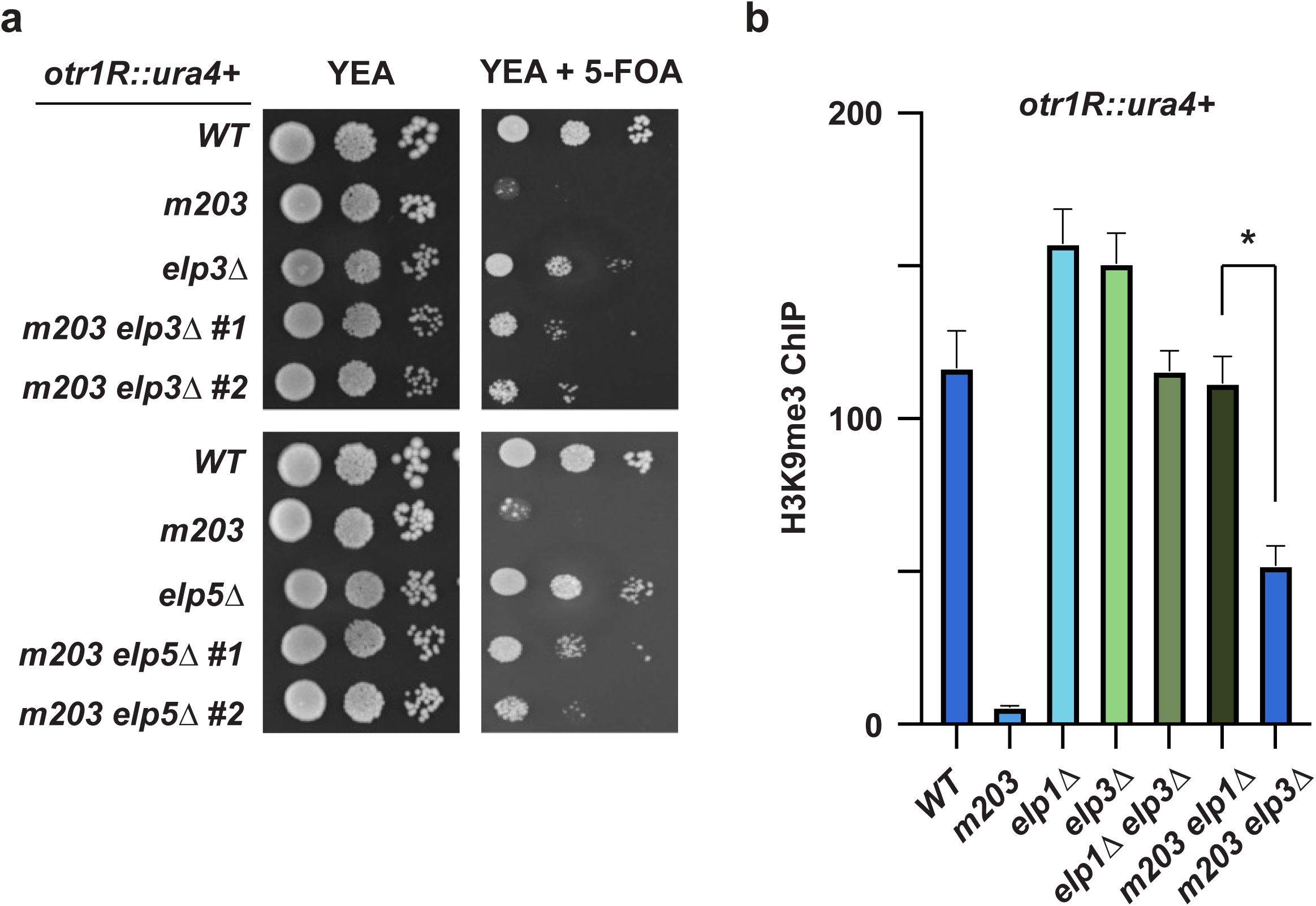
Elp1 and Elp3 have distinct contribution to heterochromatin regulation. (A) Spotting assays depicting 5-FOA sensitivities of the indicated yeast strains. For the *m203 elp3*Δ and *m203 elp5*Δ strains, two independent isolates are shown. (B) ChIP enrichment of heterochromatic H3K9me3 at *otr1R::ura4^+^* in the indicated yeast strains. Enrichments were normalized relative to the euchromatic *leu1* gene and a 5% input control. Error bars indicate standard deviation based on *n* ≥ 2. Asterix (*) denotes statistical significance as determined by Student’s t-test.

### Elp3-mediated tRNA modification is dispensable for heterochromatin loss in *m203* cells

From yeast to humans, the Elongator complex plays a well-characterized role in promoting the mcm^5^s^2^ modification of wobble uridine (U_34_) of tRNAs that recognize AA/AG-ending mRNA codons (Huang *et al*. 2005; Karlsborn *et al*. 2014b; Abbassi *et al*. 2020). This modification promotes proper codon-anticodon base pairing to help prevent ribosome stalling and co-translational polypeptide misfolding (Laguesse *et al*. 2015; Nedialkova and Leidel 2015; Abbassi *et al*. 2020). Since AA/AG-ending codons are pervasive throughout the transcriptome (Bauer *et al*. 2012; Goffena *et al*. 2018), it is thought that the loss of tRNA modification function of the Elongator complex would lead to pleiotropic downstream effects, including transcriptional dysregulation, altered exocytosis, and cytoskeletal reorganization (Esberg *et al*. 2006; Svejstrup 2007; Johansson *et al*. 2018).

To determine whether defects in mcm^5^s^2^ tRNA modifications could explain the epigenetic role of Elp1 in *m203* cells, we introduced substitution mutations within the Elp3 catalytic subunit of the Elongator complex to specifically prevent tRNA modifications while keeping the *elp1* gene intact. First, we aligned the Elp3 protein sequence from *M. jannaschii*, *S. pombe*, and *S. cerevisiae* to identify a homologous iron-sulfur cluster (Fe_4_S_4_) because it is known to be required for the mcm^5^s^2^U_34_ modification function of the Elongator complex (Dauden *et al*. 2019) (Fig. 6a). The cluster was previously experimentally verified for Elp3 from *M. jannaschii* and *S. cerevisiae* (Paraskevopoulou *et al*. 2006; Abbassi *et al*. 2024). We identified a pair of conserved cysteine residues at Elp3 positions 106 and 109 in *S. pombe* (Fig. 6a). Next, using CRISPR-Cas9, we substituted those cysteines for alanines to generate an *elp3-*(*C106A,C109A*) mutant strain that we called *elp3-CC*. We first confirmed that Elp1 protein was expressed in *elp3-CC* mutant cells (Fig. S4a). Next, we performed mass spectrometry of total tRNAs to confirm that *elp3-CC* cells completely lost mcm^5^s^2^ tRNA modifications (Fig. 6b), which is also true for tRNAs from *elp1*Δ and *elp3*Δ cells (Huang *et al*. 2005; Karlsborn *et al*. 2014b). Due to the lack of tRNA modification capability, the *elp3-CC* mutant was hypersensitive to H_2_O_2_-mediated oxidative stress (Fig. S4b), as previously reported for the *elp1*Δ and *elp3*Δ mutants (FernÁndez-VÁzquez *et al*. 2013). These results demonstrate that our Elp3-CC mutant completely ablates mcm^5^s^2^ tRNA modifications while keeping Elp1 protein expression intact.

**Fig 6.**
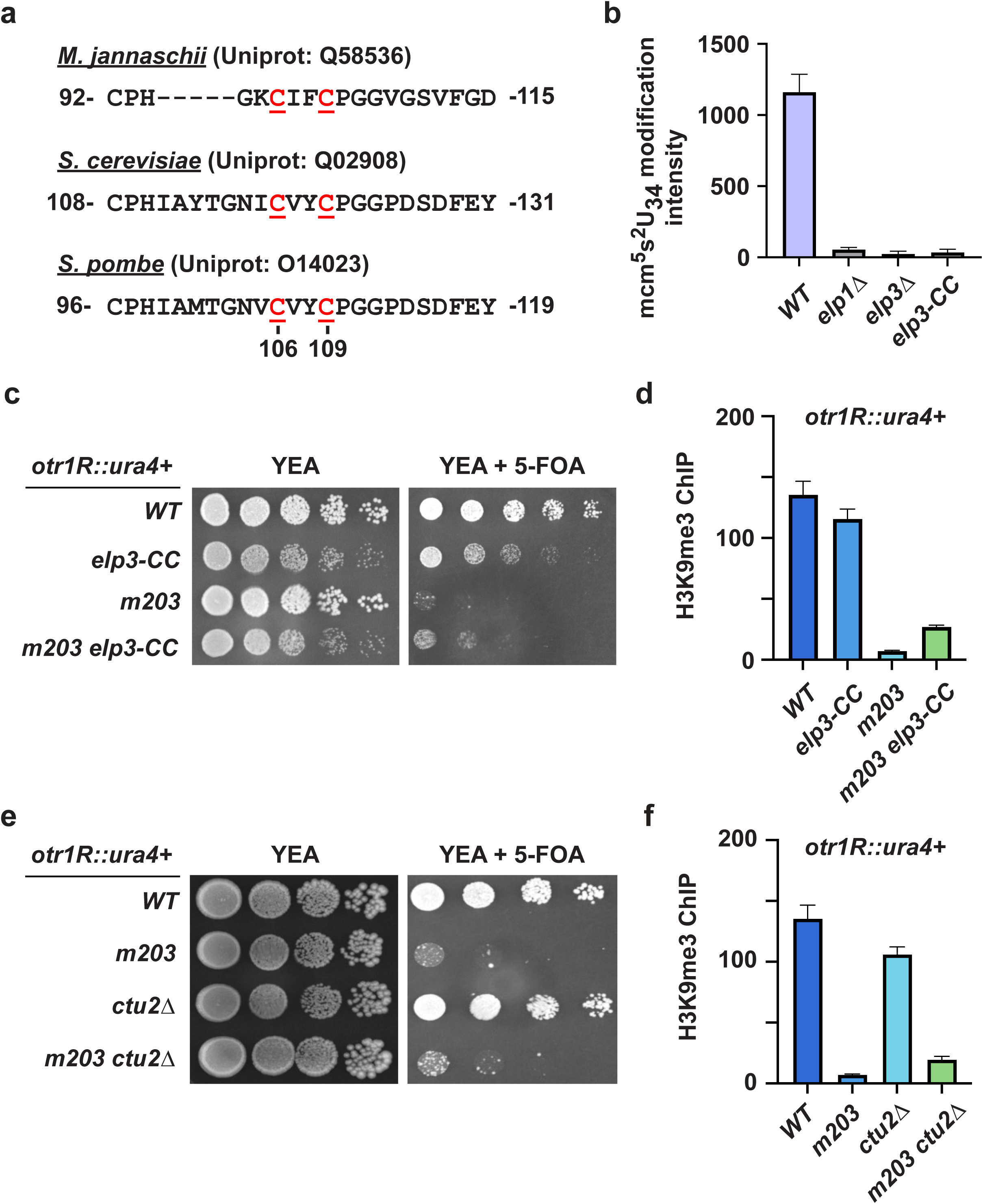
The loss of mcm^5^s^2^U_34_ tRNA modifications does not restore heterochromatic repression of *otr1R::ura4^+^* in *m203* cells. (A) Alignment of homologous Elp3 protein sequences from the thermophilic archea *Methanocaldococcus jannaschii*, budding yeast *Saccharomyces cerevisiae*, and fission yeast *Schizosaccharomyces pombe*. A partial view of the alignment that is focused on the conserved cysteine residues of the iron-sulfur cluster (underlined in red) is shown. Cysteine residues at positions 106 and 109 of *S. pombe* Elp3 were substituted for alanines to generate the Elp3-CC mutant. (B) Peak area of detected 5-methoxycarbonylmethyl-2-thiouridine (mcm^5^s^2^U) in the total tRNAs that were derived from the indicated yeast strains. (C) Spotting assays depicting 5-FOA sensitivities of the indicated yeast strains. (D) ChIP enrichment of heterochromatic H3K9me3 at *otr1R::ura4^+^*in the indicated yeast strains. (E) Spotting assays depicting 5-FOA sensitivities of *WT*, *m203*, *ctu2*Δ, and *m203 ctu2*Δ yeast strains. (F) ChIP enrichment of heterochromatic H3K9me3 at *otr1R::ura4^+^* for the yeast strains shown in panel E. ChIP enrichments were normalized relative to the euchromatic *leu1* gene and a 5% input control. Error bars indicate standard deviation based on *n* ≥ 2. For panels D and F, the enrichments for *WT* and *m203* strains are reused because they were performed in the same experiment.

To test whether *elp3-CC* could restore heterochromatic repression of the *otr1R::ura4+* reporter in *m203* cells, we compared single and double mutants using our 5-FOA sensitivity assay. Similar to the *elp3*Δ mutant, *elp3-CC* did not strongly restore growth of *m203* cells on 5-FOA media (Fig. 6c). Validating this finding, H3K9me3 ChIP-qPCR analysis showed that *elp3-CC* only slightly restored H3K9me3 at *otr1R::ura4+* in *m203* cells (Fig. 6d). In addition to using our *elp3-CC* mutant, we also deleted the *ctu1* gene (*ctu1*Δ), which encodes for the Ctu2 tRNA thiolase that is required for adding the sulfur group (s^2^) within the mcm^5^s^2^ modification (Dewez *et al*. 2008), and found weak restoration of *otr1R::ura4+* repression in *m203* cells (Fig. 6e). There was also minor restoration of H3K9me3 at the reporter locus in cells *m203 ctu1*Δ (Fig. 6f). Altogether, these data demonstrate that mcm^5^s^2^ tRNA modifications play minor roles in the heterochromatin regulatory function of Elp1 at our pericentromeric heterochromatin reporter.

Elongator complex-mediated tRNA modifications promote translation by preventing ribosomal stalling (Nedialkova and Leidel 2015). Therefore, we tested whether partial global translational inhibition would be sufficient to restore heterochromatic repression at *otr1R::ura4*^+^ in *m203* cells. Using 20μM of cycloheximide, there was no restoration of *m203* cell growth on 5-FOA media, indicating that *otr1R::ura4*^+^ was not repressed upon pan-reduction of translation (Fig. S5). Along with our *elp3-CC* results, this suggests that the epigenetic role of Elp1 involves processes beyond mcm^5^s^2^ tRNA modifications and global translation.

### Overexpression of tRNA^Lys^_UUU_ that lack mcm^5^s^2^ modification suppresses *elp1Δ*

It was previously shown in *S. cerevisiae* and *S. pombe* yeast models that simply overexpressing tRNA^Lys^_UUU_, but not tRNA^Lys^_CUU_, in *elp3*Δ cells would suppress Elongator-associated defects in transcription, cell cycle, and stress response (Esberg *et al*. 2006; Bauer *et al*. 2012; FernÁndez-VÁzquez *et al*. 2013). In wild-type cells, the Elongator complex promotes mcm^5^s^2^ modification of U_34_ from tRNA^Lys^_UUU_ (Huang *et al*. 2005; Karlsborn *et al*. 2014b). However, tRNA^Lys^_CUU_ cannot have the same modification because the wobble nucleoside is cytosine. Nonetheless, both tRNAs are charged with the lysine amino acid, making them well suited for side-by-side comparisons of tRNAs that are normally modified, or not, by the Elongator complex (Esberg *et al*. 2006; Bauer *et al*. 2012). This tRNA overexpression strategy was previously used to suggest that the Elongator complex relies on certain tRNAs to regulate diverse processes (Esberg *et al*. 2006; Bauer *et al*. 2012). We aimed to test whether this also applies to the epigenetic function of Elp1.

We cloned the *SPBTRNALYS.06* and *SPCTRNALYS.11* genes, which encode for tRNA^Lys^_UUU_ and tRNA^Lys^_CUU_, respectively, along with 500 bp of their upstream and downstream regions (to include promoter and terminator elements), into high-copy yeast plasmids that contained the *NatMX* selectable marker gene (allowing growth on media with the nourseothricin antibiotic). We found that overexpression of tRNA^Lys^_UUU_ enhanced the colony sizes of *m203 elp1*Δ cells compared to vector-only and tRNA^Lys^_CUU_ overexpression controls (Fig. S6), agreeing with previous findings that tRNA^Lys^_UUU_ overexpression can rescue growth defects in *S. cerevisiae* and *S. pombe* Elongator mutant cells (Esberg *et al*. 2006; Villahermosa and Fleck 2017). We next tested whether tRNA^Lys^_UUU_ overexpression in *m203 elp1*Δ cells can revert our *otr1R::ura4^+^* reporter from a repressed to an expressed state. We found that tRNA^Lys^_UUU_ overexpression re-sensitized *m203 elp1*Δ cells to 5-FOA (Fig. 7a). In contrast, the empty vector or overexpression of tRNA^Lys^_CUU_ showed far less suppressive effects (Fig. 7a). By H3K9me3 ChIP analyses, we also found that overexpression of tRNA^Lys^_UUU_, but not tRNA^Lys^_CUU_, in *m203 elp1*Δ cells dramatically reduced heterochromatic H3K9me3 at the *otr1R::ura4^+^* locus (Fig. 7b). Taken together, these data indicate that tRNA^Lys^_UUU_ overexpression can suppress the epigenetic function of Elp1.

**Fig 7.**
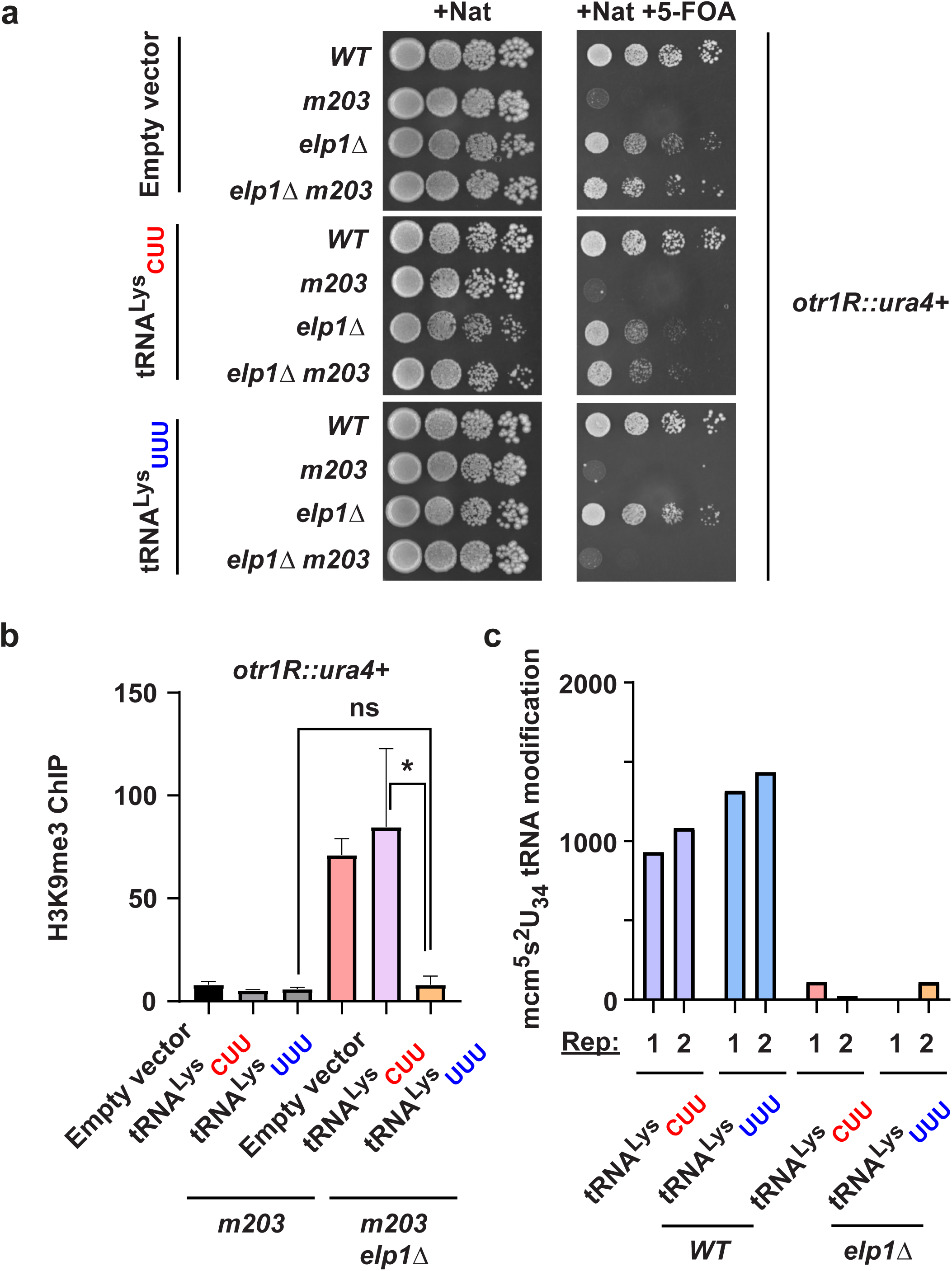
Overexpression of tRNA^Lys^_UUU_ in *m203 elp1*Δ cells reverts the *otr1R::ura4^+^*reporter back to the expressed state. (A) Spotting assays showing 5-FOA sensitivity of the indicated yeast strains which carry empty vector or expression plasmids with tRNA^Lys^_CUU_ (*SPCTRNALYS.11*) or tRNA^Lys^_UUU_ (*SPBTRNALYS.06*). All yeast strains were grown on media containing nourseothricin antibiotic (Nat) to select for cells with the plasmids. (B) ChIP enrichment of heterochromatic H3K9me3 at *otr1R::ura4^+^* in *m203* or *m203 elp1*Δ cells that were shown in panel A. (C) Peak area of detected 5-methoxycarbonylmethyl-2-thiouridine (mcm^5^s^2^U) in the total tRNAs that were derived from *WT* or *elp1*Δ yeast strains. Depicted are the peak area intensities from two biological replicates, per yeast strain. ChIP enrichments were normalized relative to the euchromatic *leu1* gene and a 5% input control. Error bars indicate standard deviation based on *n* ≥ 2. Asterix (*) denotes statistical significance as determined by Student’s t-test.

The tRNA overexpression experiments described above appeared to contradict our earlier conclusions using the *elp3-CC* mutant. Therefore, we wondered whether *elp1*Δ cells could regain mcm^5^s^2^ tRNA modifications when they carried the tRNA^Lys^_UUU_ plasmid. After isolating total tRNAs and subjecting them to mass spectrometry analyses, we failed to detect mcm^5^s^2^ modifications in cells lacking *elp1* (Fig. 7c), indicating that the overexpressed tRNAs do not have the modification. Our finding agrees with a previous study in *S. cerevisiae* that overexpressed tRNAs in Elongator mutant cells fail to regain mcm^5^s^2^ (Esberg *et al*. 2006). We conclude that Elp1-mediated loss of heterochromatic silencing in *m203* cells is dependent on Elongator complex-modifiable tRNA^Lys^_UUU_ but not on the mcm^5^s^2^ modification function, *per se*.

## Discussion

Our study reveals that the highly conserved Elp1 protein may possess a novel function in negatively regulating RNAi-dependent heterochromatin. Surprisingly, this epigenetic role of Elp1 may involve mechanisms that are independent of other Elongator complex subunits and of the mcm^5^s^2^ tRNA modification, representing the first instance to our knowledge that Elp1 might be capable of an Elongator-independent function. This work expands our understanding of Elp1 functional capabilities beyond the current paradigms of tRNA modifications (Dauden *et al*. 2019) and microtubule assembly (Planelles-Herrero *et al*. 2022; Planelles-Herrero *et al*. 2025), both of which require all Elongator complex subunits and their ability to promote mcm^5^s^2^ tRNA modifications.

In this study, an epigenetic role for Elp1 was revealed in the context of a tester *S. pombe* strain called *m203*. Because this unique strain disrupts the RNAi pathway while keeping transcription intact at the RNAi-dependent heterochromatin reporter locus *otr1R::ura4^+^* (Kato *et al*. 2005), it provided an elegant system to identify novel factors that could mediate the RNAi disruption. Interestingly, loss of *elp1* not only restored heterochromatic repression in *m203* cells but also in *rpb2-N44F* cells (Fig. S2b), suggesting that there might be several contexts where Elp1 could have an epigenetic role. Supporting this idea, a recent genome-wide study reported that Elp1 can regulate heterochromatic repression at centromeres, subtelomeres, and the mating-type locus (Muhammad *et al*. 2024). Future work will be required to define the mechanism(s) by which Elp1 regulates RNAi and heterochromatin.

Intriguingly, the other Elongator complex subunits Elp2/3/4 have been reported to promote RNAi-dependent heterochromatin (Muhammad *et al*. 2024). Additionally, an earlier study also reported that Elp3 positively regulates heterochromatin at the *otr1R::ura4^+^* reporter (Bauer *et al*. 2012), which we used in our study. It was suggested that Elp3 regulates heterochromatin by facilitating the translation of RNAi factors, including Ago1, via its mcm^5^s^2^ tRNA modification function (Bauer *et al*. 2012). In our own studies, we did notice that *elp3*Δ and *elp5*Δ mutants had small but noticeable effects on heterochromatic repression (Figs. 5a, b). Nonetheless, we found that the heterochromatic phenotypes of *elp1*Δ mutants were more prominent. We propose that Elp1 may regulate heterochromatin via Elongator complex-dependent and -independent mechanisms. Future studies employing separation-of-function constructs, such as our *elp3-CC* mutant, will be needed to delineate these mechanisms.

Overexpression of tRNA_UUU_ robustly alleviates heterochromatic repression in *m203 elp1*Δ cells but total inhibition of mcm^5^s^2^ tRNA modifications, by our *elp3*Δ or *elp3-CC* mutant, in *m203* cells only had minor effects (Figs. 5a and 6c). This differs from the conclusions of other investigations that focused on other processes, including transcription, exocytosis, or stress response, where genetic inactivation of the Elongator complex and tRNA_UUU_ overexpression yielded opposite, but complementary, phenotypes (Esberg *et al*. 2006; Bauer *et al*. 2012; FernÁndez-VÁzquez *et al*. 2013). Our results could be explained through Elongator-dependent and -independent mechanisms of heterochromatin regulation by Elp1. Key to this hypothesis is that we and others have found that the overexpressed tRNAs do not become mcm^5^s^2^-modified in Elongator mutants (Fig. 7d) (Esberg *et al*. 2006). High levels of tRNA_UUU_ lacking mcm^5^s^2^ may help correct defects in translational decoding of AAA_Lys_ and AAG_Lys_ codons (Nedialkova and Leidel 2015). However, tRNA overexpression could also lead to widespread reprogramming within cells in poorly understood ways (Pavon-Eternod *et al*. 2013). It is possible that tRNA_UUU_ overexpression perturbed our heterochromatin system through pleiotropic mechanisms. Future exploration of molecular consequences due to overexpression of different tRNAs may provide clues into pathways that Elp1 could leverage for its epigenetic role.

The homolog of *S. pombe* Elp1 in humans is ELP1 (previously known as IKAP). Intronic mutations that prevent proper splicing of the *ELP1* mRNA cause Riley-Day syndrome, a recessive autosomal condition characterized by progressive neurodevelopmental and neurodegenerative aberrations (Anderson *et al*. 2001; Slaugenhaupt *et al*. 2001). Although it is clear that *ELP1* is critically important for neuronal health, the molecular basis behind how human ELP1 functions within neurons remains elusive. It is known that mis-splicing of ELP1 results in reduced ELP1 protein expression and lowered mcm^5^s^2^ tRNA modifications (Karlsborn *et al*. 2014a), however, it is unclear the extent to which these outcomes cause disorders. Furthermore, there are no reports of mutations in the other Elongator complex subunits (ELP2/3/4/5/6) that cause Riley-Day syndrome, suggesting that neuronal ELP1 might be important for other Elongator complex-independent processes or that subtleties in the process of Elongator-mediated tRNA modification help distinguish Elongator-associated diseases and disorders (Gaik *et al*. 2023). Future studies using yeast and animal models, combined with separation-of-function tools, will be important for investigating the underexplored Elongator complex–independent roles of Elp1/ELP1 and its impact on human health.

---

**Fig S1.**
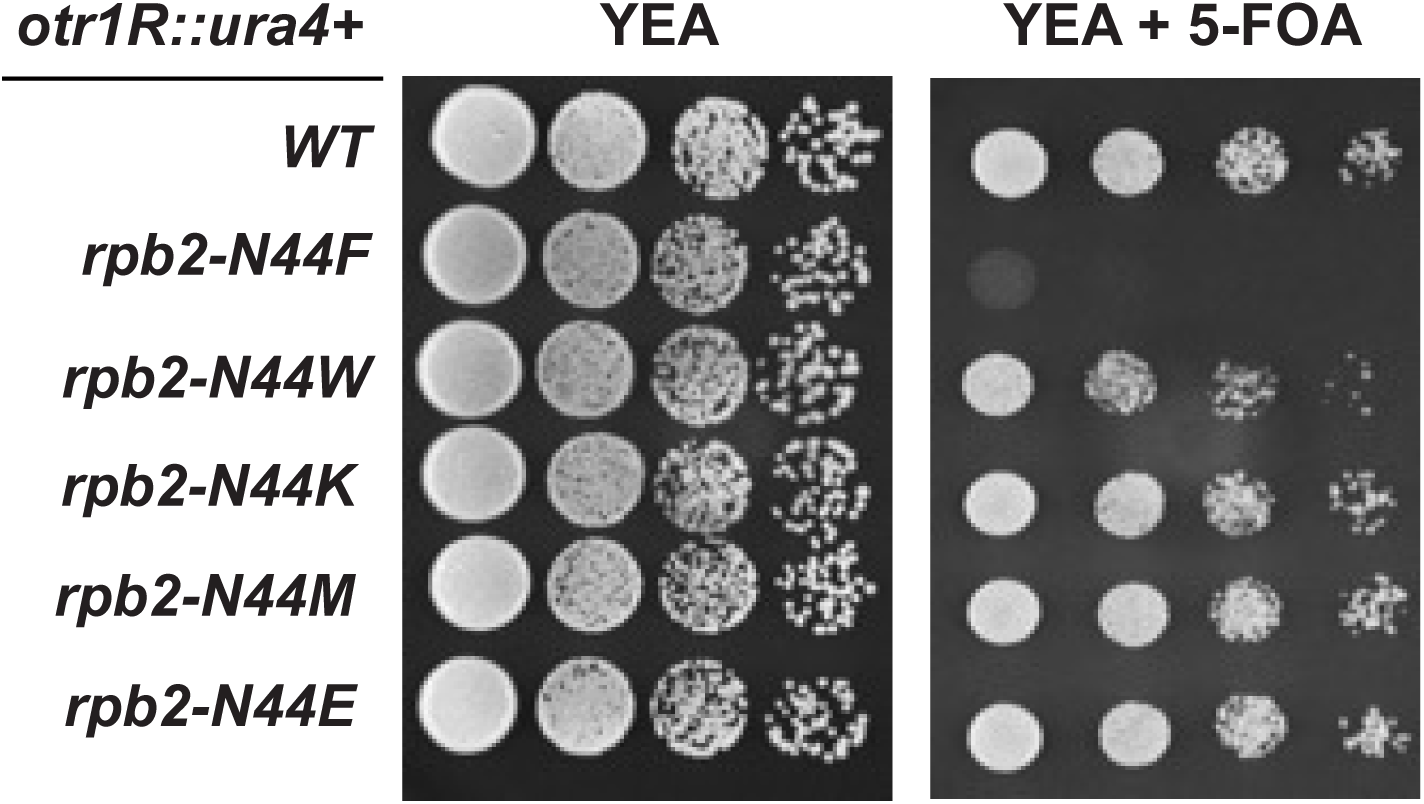
Not all substitutions of Rpb2-N44 in yeast with the *otr1R::ura4^+^* reporter promote 5-FOA sensitivity. Spotting assay showing 5-FOA sensitivity of the indicated yeast strains. Here, only substitution of Rpb2-N44 with phenylalanine (F) led to 5-FOA hypersensitivity.

**Fig S2.**
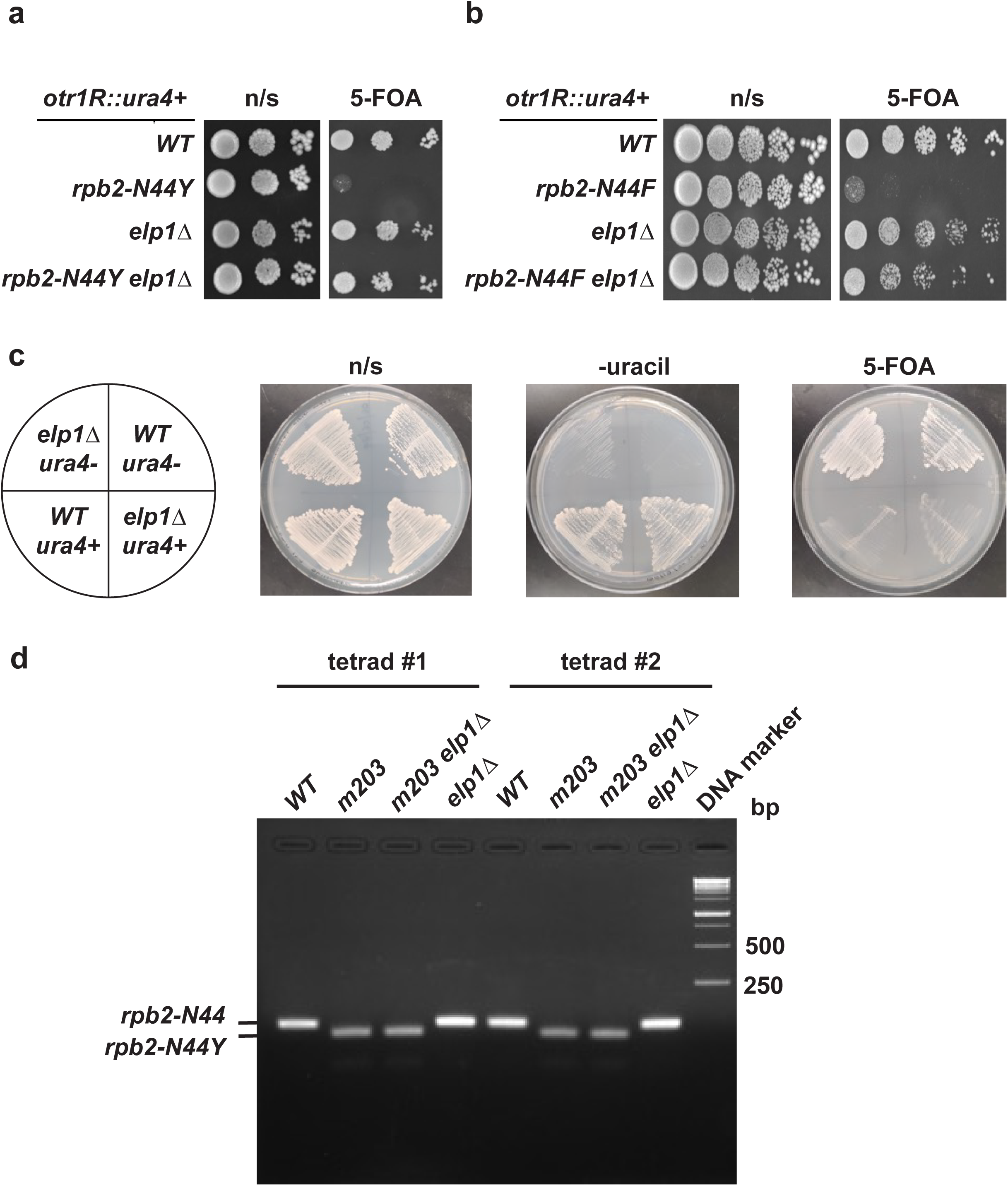
*Elp1* genetically interact with *rpb2-N44Y* (*m203*) and *rpb2-N44F* variant alleles. (A and B) Spotting assay showing 5-FOA sensitivity of the indicated yeast strains. (C) Cell spreading assay of the yeast strains that are indicated in the illustration to the left. The strains either expressed *ura4* from its endogenous euchromatic locus (*ura4+*) or completely lacked the *ura4* gene (*ura4-*). Yeast strains were spread onto non-selective (n/s) YEA media plate, plate media lacking uracil, or plate media containing 5-FOA. (D) *m203* and *elp1*Δ yeast strains were crossed to produce tetrads that contained sibling yeast progeny, and dCAPS was performed to delineate the *rpb2* alleles (*rpb2-N44* and *rpb2-N44Y*) within each progeny. Shown are dCAPS results for yeast progeny from two representative tetrads. A DNA marker is also shown for size reference, in base-pairs (bp).

**Fig S3.**
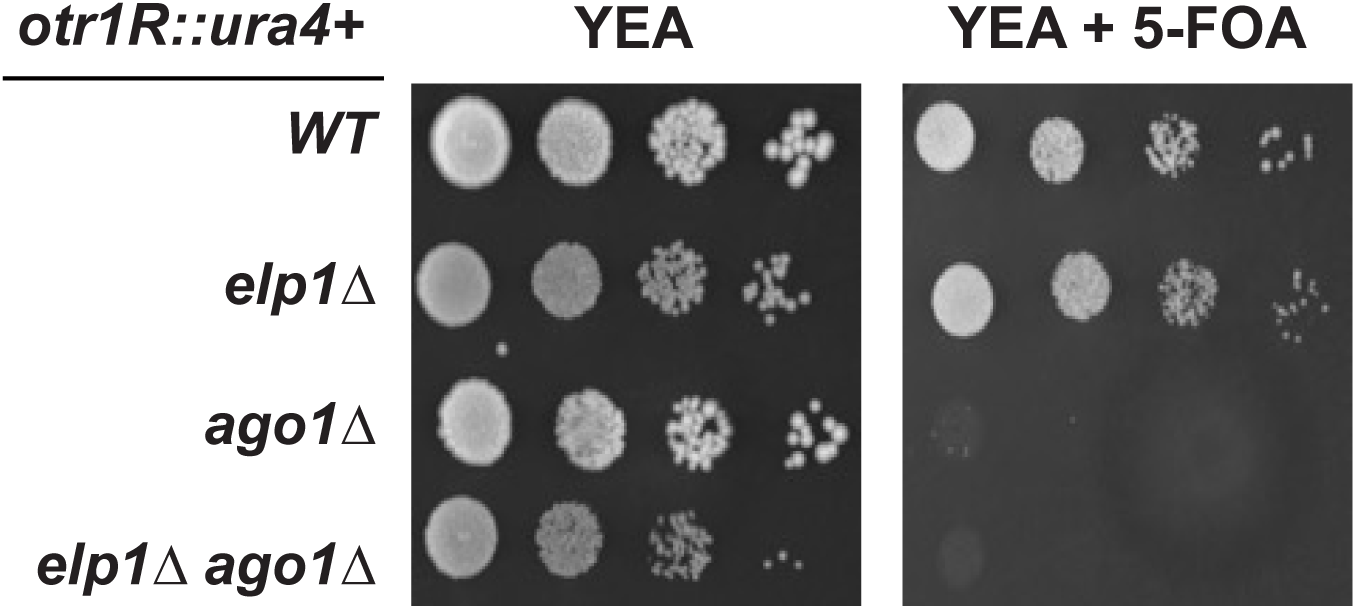
Loss of *elp1* does not restore 5-FOA hypersensitivity in *ago1*Δ cells with the *otr1R::ura4^+^* reporter. Spotting assay showing 5-FOA sensitivity of the indicated yeast strains.

**Fig S4.**
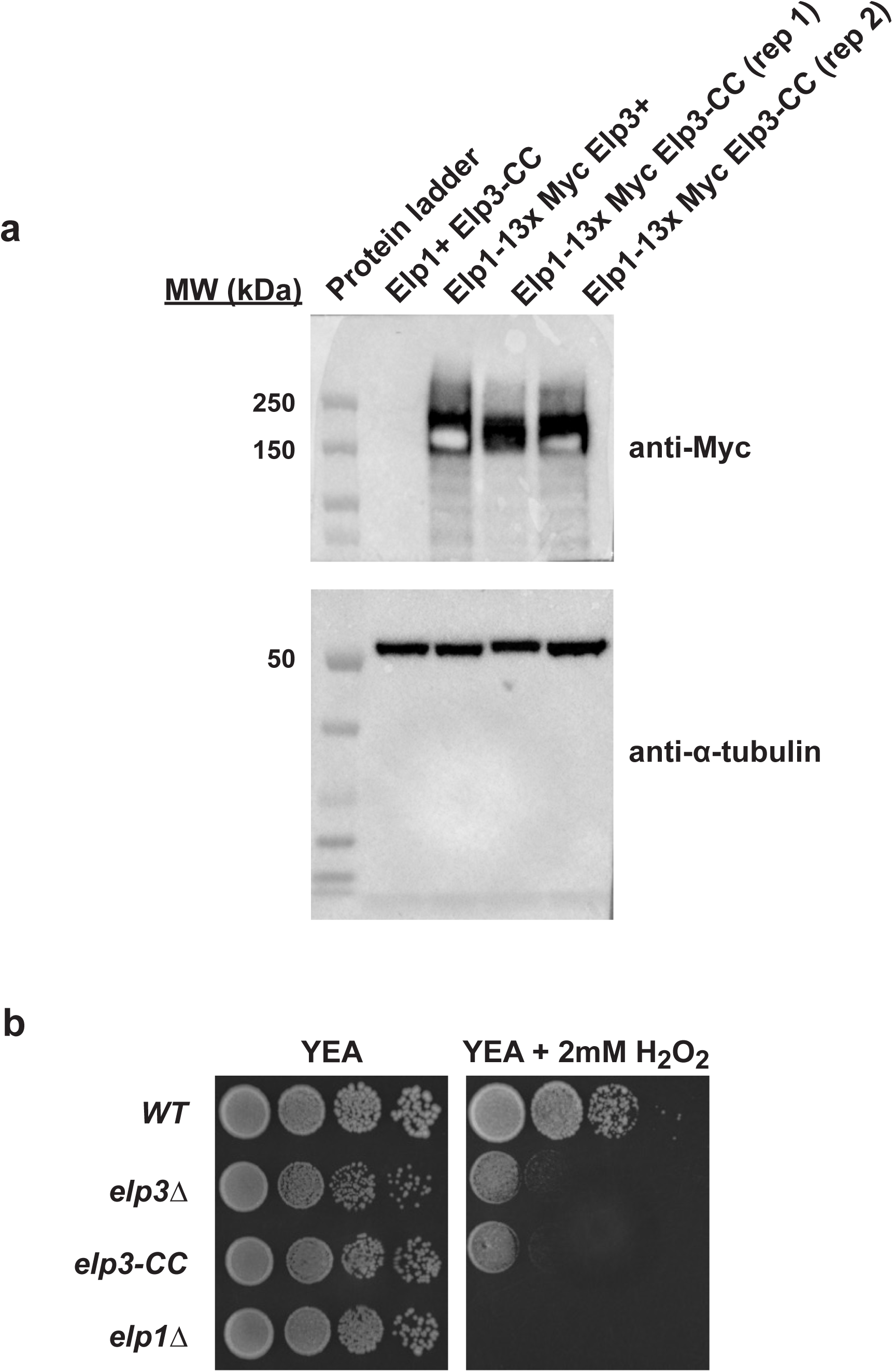
Elp1 protein is expressed in the *elp3-CC* yeast mutant. (A) Western blot analysis of untagged or 13x Myc-tagged Elp1 in yeast strains that either have wild-type *elp3* (*elp3+*) or *elp3-CC*. α-Tubulin was probed as a loading control. Also shown is the protein ladder for size reference, in kilodaltons (kDa). (B) Spotting assay showing sensitivity of the indicated yeast strains to 2 mM hydrogen peroxide (H_2_O_2_).

**Fig S5.**
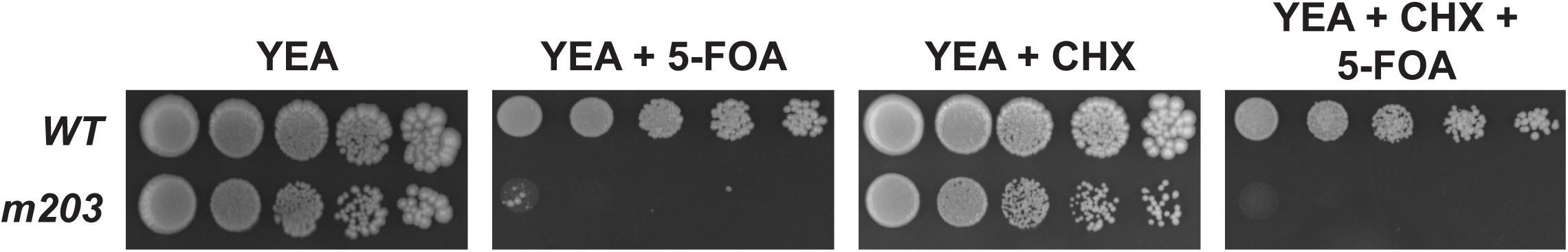
Translational inhibition with cycloheximide (CHX) does not suppress 5-FOA hypersensitivity of *m203* cells with the *otr1R::ura4^+^* reporter. Spotting assay showing sensitivity to 5-FOA, in the absence or presence of 20 mM CHX, of *WT* or *m203* yeast cells.

**Fig S6.**
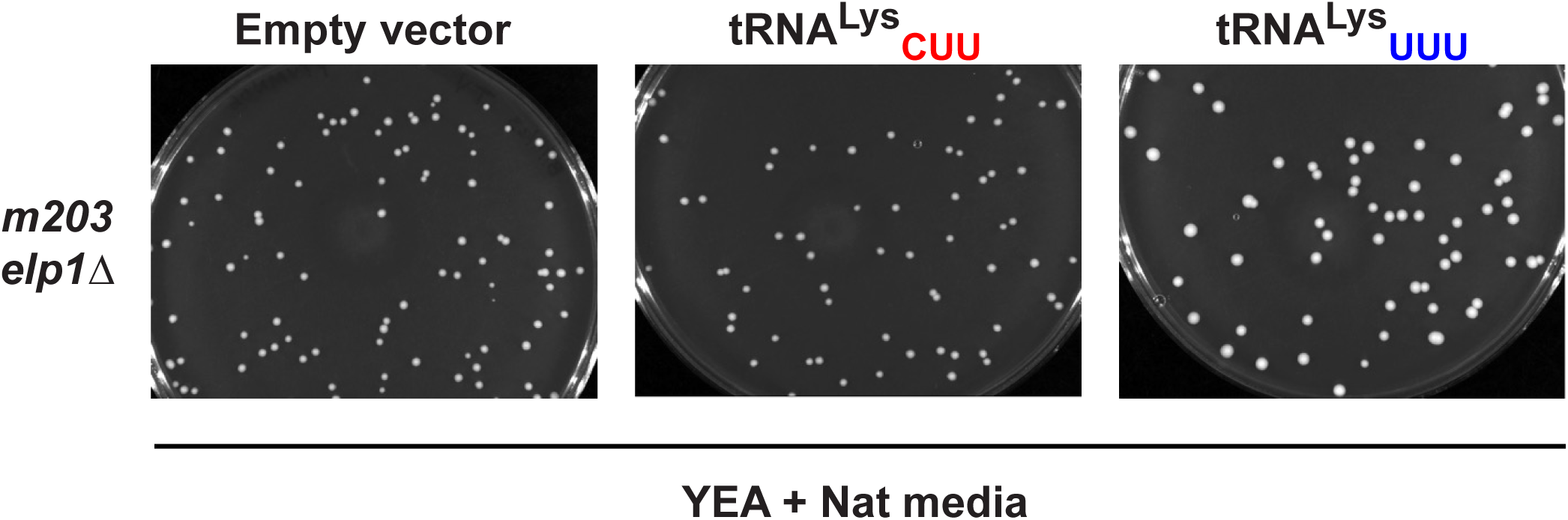
Plasmid-derived expression of tRNA^Lys^_UUU_ enhances yeast colony size of *m203 elp1*Δ cells. Selective YEA media plates (containing nourseothricin, Nat) with single colony spreads of *m203 elp1*Δ yeast strains that contain empty or tRNA expression vectors. *m203 elp1*Δ yeast colonies typically grow smaller compared to *WT*. *m203 elp1*Δ cells that overexpress tRNA^Lys^_UUU_ produce larger colonies that approximately resemble *WT* sizes.

## Data availability

The strains and plasmids are available upon reasonable request to the corresponding author.

## Acknowledgements

All data shown here were generated by the Vo lab. We would like to thank Dr. Shiv Grewal for sharing support, yeast strains, and protocols. We thank Dr. Yota Murakami for sharing the *m203* yeast strain from his lab. We are also grateful to Drs. David Arnosti, Sebastian Glatt, and Jesper Svejstrup for their helpful scientific suggestions about this work, and Dr. Benjamin Orlando for technical help with interpreting Elp3 protein domain data. We acknowledge Novogene for their service in producing the small RNA-seq libraries and raw sequencing results. We acknowledge ArrayStar, Inc. for their service in purifying tRNAs and detecting nucleoside modifications via LC-MS. We acknowledge the MSU Bioinformatics core facility for their help in the small RNA-seq bioinformatic analyses. Lastly, we would like to thank all current and former members of the Vo lab for their helpful suggestions during this work.

## Author contributions

T.V.V. and M.B.N. conceived and designed the project. M.B.N., M.E.P., C.T.L., J.M.F., K.S.D., and T.V.V. performed experiments. M.B.N. and T.V.V. performed the majority of the experiments and analyses. M.E.P. and C.T.L. performed replicating and other support experiments. J.M.F. and K.S.D. performed experiments for Figure 6C. M.B.N. prepared raw figures and tables, and wrote a draft of the Materials and Methods. T.V.V. prepared the remainder of the manuscript with input from all authors.

